# Strength of functional signature correlates with effect size in autism

**DOI:** 10.1101/043422

**Authors:** Sara Ballouz, Jesse Gillis

**Affiliations:** The Stanley Institute for Cognitive Genomics, Cold Spring Harbor Laboratory, Cold Spring Harbor, NY, 11724, USA

**Author notes:** Corresponding author: Dr J Gillis, The Stanley Institute for Cognitive Genomics, Cold Spring Harbor Laboratory, Cold Spring Harbor 11724, NY, USA.

**Keywords:** autism spectrum disorder, rare variation, common variation, loss-of-function, recurrence, effect sizes, functional enrichment, gene candidate score, meta-analysis

## Abstract

**Background:** Disagreements over genetic signatures associated with disease have been particularly prominent in the field of psychiatric genetics, creating a sharp divide between disease burdens attributed to common and rare variation, with study designs independently targeting each. Meta-analysis within each of these study designs is routine, whether using raw data or summary statistics, but combining results across study designs is atypical. However, tests of functional convergence are used across all study designs, where candidate gene sets are assessed for overlaps with previously known properties. This suggests one possible avenue for combining not study data, but the functional conclusions that they reach.

**Method:** In this work, we test for functional convergence in autism spectrum disorder (ASD) across different study types, and specifically whether the degree to which a gene is implicated in autism is correlated with the degree to which it drives functional convergence. Because different study designs are distinguishable by their differences in effect size, this also provides a unified means of incorporating the impact of study design into the analysis of convergence.

**Results:** We detected remarkably significant positive trends in aggregate (p < 2.2e-16) with 14 individually significant properties (FDR<0.01), many in areas researchers have targeted based on different reasoning, such as the fragile X mental retardation protein *(FMRP)* interactor enrichment (FDR 0.003). We are also able to detect novel technical effects and we see that network enrichment from protein-protein interaction data is heavily confounded with study design, arising readily in control data.

**Conclusions:** We see a convergent functional signal for a subset of known and novel functions in ASD from all sources of genetic variation. Meta-analytic approaches explicitly accounting for different study designs can be adapted to other diseases to discover novel functional associations and increase statistical power.

## Background

Over the last decade, enormous progress has been made in characterizing sources of DNA variation contributing to disease. Most of this progress has been enabled by study designs which are carefully tailored to exploit technologies targeting particular classes of variation. Researchers have used chromosomal analysis arrays [1-4], genotyping arrays [5-8], whole-exome sequencing (WES)[9-14], and whole genome sequencing (WGS)[15, 16], to identify risk loci and alleles. The results from these studies cannot be naively compared; common variants are limited to regions of the genome with known variation (a SNP is known) but only reach significance with large numbers, while rare or ultra-rare variants are conditioned on not being in this list of common variants. Trio and quad studies are used mainly in WES and WGS study designs, while large case and control cohorts are required for signals in genome-wide association studies (GWAS). Thus for each study design, we are asking distinct questions that relate to the population prevalence, disease mechanism, burden and risk.

Within each study, however, it is commonplace to look to overlapping functional properties of candidate disease genes to find the biologically meaningful signal among the positive results. Candidate genes are prioritized based on enrichment analyses in pathways related to the phenotype (e.g., neuronal activity regulation) or some other disease feature shared by the genes (e.g., expression in the brain). If these methods return no significant results, more complex methods are performed to extract common features from the disease gene set [17], such as co-regulatory module detection from co-expression networks [18] or binding from protein-protein interaction (PPI) networks [19]. Regardless of the study design, the analysis with respect to functional convergence follows a similar (and largely separable) design: genes selected as hits are tested for the presence of some joint signature with the null provided by genes which are not hits. By the same logic that suggests testing hits for functional convergence relative to the background, we hypothesize that sets of genes which are "strong" hits will show more functional convergence than those which are "weak" hits.

We suggest that the degree of functional convergence may by hypothesized to vary (monotonically) with the degree to which genes are causal for the disease. Genes only weakly causal, whether due to high false positive rates in the study design or low effect sizes, are not strongly implicated as sharing a joint role by their co-occurrence as disease-related. For instance, disease candidates from GWAS have low relative risks (and therefore low effect sizes) as they are inherited common variation in the population. On the other hand, *de novo* mutations are a form of genetic variation which evolutionary forces have had little time to act upon [20] (e.g., unless embryonically lethal), and are of high risk (and high effect sizes). Studies also suffer from type I errors (false positives), and this too should be reflected in an aggregate disease signal of the candidate genes, as quantified by their common functional properties. A set of genes with *de novo* mutations will show a strong aggregate disease signal, while we might expect a weaker signal from the gene candidates from GWAS [21]. Measuring their 'functional convergence', as determined by a gene set enrichment test or network analysis, we can thus exploit our knowledge of gene candidates' effect sizes and false positive rates. For a true disease property, we expect the correlation between gene set effect size and functional convergence to be strong, and for a weak or artifactual property, we expect no significant correlation.

We propose to test this hypothesis by running a meta-analytic study on autism spectrum disorder (ASD [MIM 209850]) candidates across numerous genetic studies and over a wide range of gene properties and functions. ASD is a neurodevelopmental disease commonly characterized by behavioral traits such as poor social and communication skills [22]. In more severe cases, ASD is comorbid with mild to severe intellectual disability, facial and cranial dysmorphology and gastrointestinal disorders. Perhaps because of grouping these multiple and sometimes distinct phenotypes into one disorder, and the complexity of behavior as a trait, understanding the genetic architecture of this cognitive disease has been non-trivial [23]. The genetic component of ASD is estimated to be 50-60% [24], however there are still a substantial number of cases where the underlying genetic factors of the disease are unknown. Due to these levels of heterogeneity, multiple studies and study designs have been used to determine the underlying genetics which we make use of here. Taking these different studies, we construct several disease gene candidate collections, each containing genes of similar levels of risk, as determined by their odds ratios and relative risks. On every gene collection, we run a number of analyses, calculating the functional convergence using standard enrichment methods, and more complex network analysis enrichments. By exploiting trends in targeted genetic variation and their known effect sizes, we demonstrate it is possible to discriminate biologically convergent signals from likely technical artifacts at a very fine resolution. The disease properties with strong trend signals are largely consistent with the known literature on ASD (e.g., FMRP interactor enrichment) but we also see a few otherwise interesting properties as unlikely to be disease specific. Particularly protein-protein interaction networks and some co-expression networks, which extract artifactual signals from the study design, show signals in control data using that study design. Our focus here is on autism due to our interest in the disorder, its well-powered data, and also its phenotypic and genetic heterogeneity.

## Methods

### Study design

An overview of our study design and method is shown in **Fig 1**. Briefly, we start by characterizing the ASD gene sets collected for this analysis. Each study's results were collapsed individually into a set of genes, with an estimated average effect size for that candidate set (**Fig 1A**). We calculate a functional effect (e.g., statistical overlaps with known functions, **Fig 1B**) for disease-specific and more general gene properties. We then calculate the correlation of these functional convergences with the estimated effect size of that variant class (**Fig 1C**). More specifically, we test to see if the set of genes with high effect sizes have strong relative functional convergences as measured by a functional enrichment of some disease property across them, and those with low effect sizes, have weaker functional signals. We apply this test to numerous functional properties on candidate gene sets from a variety of study designs. Functions with positive correlations (positive trends) we believe will show signatures that are likely associated with autism and can be used for further functional characterization of the disease. Throughout our work we refer to the "effect size" as the disease burden or risk of a gene candidate (or the average of such values within a gene set), and the "functional convergence" as the significance of a functional test for a disease gene set after controlling for the set size.

**Fig 1.**
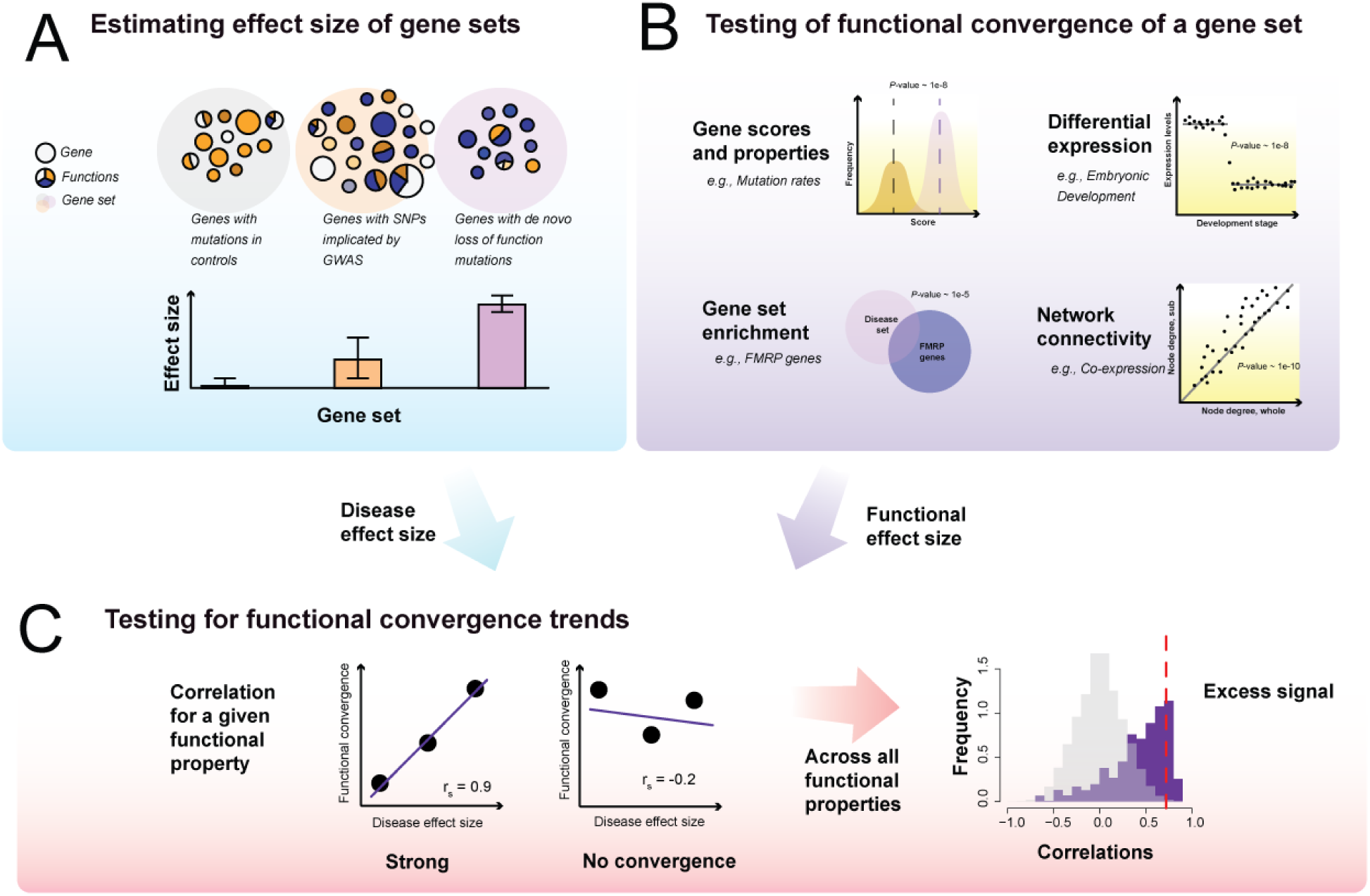
Schematic of functional convergence trend calculation. (A) Starting with disease gene set collections, we rank each by the average effect size of the genes within that set. (B) We then run 'functional tests' on these genes sets and calculate a functional convergence for each. (C) Then, using the ranking of the disease gene sets, we measure the functional convergence signature - the correlation of the trend line of the functional convergences versus the rank.

### Study data

#### Disease gene candidate sets

We first collected candidate disease gene sets from available autism studies. We selected the largest study of whole-exome sequencing (WES) of families from the Simons Simplex Collection (SSC)[25]. We defined different sets of genes from over 2000 gene candidates, splitting into recurrent (at least 2 probands having the mutation) and non-recurrent mutations, according to mutation type (loss-of-function, missense and silent mutations). We selected copy number variant (CNV) data also from the individuals in the SSC [26], and parsed it into similar sets. We then used the CNVs as parsed by Gilman et al.[3], which prioritized genes with their NETBAG algorithm. For GWAS gene sets, we generated two lists from the Psychiatric Genomics Consortium (PGC) study on autism and related psychiatric disorders [23]: one from the reported gene list and a second list of all adjacent genes as listed in the GWAS NHGRI-EBI catalog [6]. For our control and test gene sets, we took all the GWAs data in the GWAs catalog [6], totaling over 1,396 traits across 2,066 studies. For each trait, we created gene lists with the reported genes. We conditioned on traits with at least 27 genes, which left us with approximately 200 traits. Our negative control sets included using the genes with mutations in the unaffected siblings of the probands from the SSC studies. Overall, we had 11 gene sets for the main autism analysis, and 148 trait gene sets from GWAS.

### Gene functional annotation data

#### Co-expression networks

The majority of recent studies used co-expression networks from BrainSpan to illustrate network convergence among disease genes of ASD (e.g., see [27-29]). In a similar fashion, we generated a brain specific network from the BrainSpan RNA-seq data (578 samples). In addition to this, we generated an aggregate co-expression network from 28 brain tissue and cell specific microarray experiments (3,362 samples). For more general networks, we used our aggregate RNA-seq and microarray co-expression networks as previously described in Ballouz et al, [30]. In brief, these are the aggregates of 50 networks (1,970 samples) and 43 networks (5,134) samples respectively, across various tissues, cell types and conditions. As a comparison to the aggregate networks we recommend, we constructed and tested individual networks from single experiments that are more commonly used. This includes tissue-specific co-expression networks from the GTEx data [31] (29 tissues), and age specific co-expression networks (5 age groups). As additional tests, we took a further 227 RNA-seq expression datasets with at least 20 samples within each experiment from GEMMA [8], and have generated a further 454 individual human co-expression networks, using all annotated transcripts (30K, GENCODE [32]), and then only proteincoding genes (18K).

#### Protein-protein interaction networks

We used the human physical protein-protein interactions from BIOGRID (version 3.2.121)[33] and created a binary protein-protein interaction network, where each protein was a node and each protein-protein interaction is an edge. Because of the sparseness of the network, we extended the network by modelling indirect connections [34], taking the inverse minimum path length between two proteins as the weighted edge, with a maximum distance of 6 jumps roughly as described in Gillis et al,[35]. We repeated this for alternate PPI datasets including: I2D [36] (v 2.9), HPRD [37] (Release 9), HIPPIE [38] (v1.8), IntAct [39], the CCSB interactome database [40](HI-III v2.2), STRING [41](v 10), and PIPs [42]. A non-interacting protein-protein network was created from data from the negatome [43](v2).

#### Gene sets and collections

We considered common functional gene sets and neurological specific sets, as used in numerous studies, as gene sets to test for ASD candidate enrichment. These included the post synaptic density (HPSD) gene set [44], synapse sets [45], the synaptosome [46], chromatin remodelling set [47], fragile X mental retardation protein (FMRP) set [48], and gene essentiality [49]. For more standard sets, we also took the Gene Ontology [50] (GO) terms (April 2015) and KEGG pathways [51]. For each GO term, we only used evidence codes that were not inferred electronically, propagated annotations through the ontology (parent node terms inherited the genes of their leaf node terms). To minimize redundancy from GO, we restricted our enrichment analyses to GO terms groups with sizes between 20 to 1000 genes. While these GO terms and KEGG groups are used in the enrichment analyses (with the full multiple hypothesis test correction penalty). As an extension to the original study, we collected alternate gene property sets for more functional enrichment tests. For this we used all the collections from MSigDB [7] (gene sets H, C1-C7). We calculated the multifunctionality of a gene based on the number of times a gene is seen as being annotated to a function (using GO, see [52]).

#### Disease gene score sets

We used disease gene scoring methods that rank genes according to likely having damaging effects if they are mutated. This included the Residual Variation Intolerance Score (RVIS)[53], haploinsufficiency (HI) scores [54], mutational rates and constrained gene scores and probabilities (pLI) from ExAC [55].

#### Expression data

To obtain brain specific expression and differential expression information, we used three common and large sample size brain-specific transcriptomic sets. These included the Human Brain Transcriptome (GSE25159)[56], BrainSpan [39] and the Human Prefrontal Cortex transcriptome (GSE30272)[57]. We divided the samples into fetal (post-conception week - PCW) and post-birth stages, and performed a straightforward differential expression (DE) fold change analysis (averaging across these stages)[58].

### Calculating average disease effect sizes

For the 11 candidate disease and control gene sets (**Table 1**, **Fig 2A**), we ranked the set according to the overall or average "effect size" of the genes within it. For the *de novo* mutation candidates, we took the ratio of observed counts of mutations to silent mutations within the study for that class of mutations, and then the ratio of those odds between siblings to probands (as calculated in Sanders et al,[10]). To calculate this effect size for the GWAS results, we took the average odds ratios from the individual studies of each the SNP, which ranged between 1.01-1.1. For the control sets (siblings and the silent mutations), we took the effect size to be null. We then ranked the sets based on these overall effect sizes. After these calculations, we end up with three general classes: null effects (as controls), weak effects (missense and common variants) and strong effects (rarer loss-of-function and copy number variants).

**Table 1.** Disease gene sets used in the study

| Gene set | Set size<br>(genes) | Odds Ratios/<br>effect sizes | Rank |
| --- | --- | --- | --- |
| **WES results.** |  |  |  |
| *Varying effect sizes depending on mutation and recurrence.* |  |  |  |
| *Odds ratio is calculated as in Sanders et al,[10] (ratio of observed counts of mutations to silent mutations, and then ratio of those odds between siblings to probands).* |  |  |  |
| *De novo* loss-of-function, recurrent | 27 | 4.1 | 11 |
| *De novo* CNVs | 72 | 3.95 | 10 |
| *De novo* missense, recurrent | 153 | 1.6 | 9 |
| *De novo* loss-of-function | 341 | 1.5 | 8 |
| *De novo* missense | 1339 | 1.06 | 5 |
| **GWAS. Multiple hit models, low effect size per gene. Odds ratios between 1 and 1.1** |  |  |  |
| GWAS, reported genes | 49 | 1.08 | 7 |
| GWAS, adjacent genes to SNP | 116 | 1.08 | 6 |
| **Control sets, no effect** |  |  |  |
| *De novo* silent | 590 | NA | 2.5 |
| **Control groups, no disease** |  |  |  |
| *De novo* loss-of-function (sibling controls) | 174 | NA | 2.5 |
| *De novo* missense (sibling controls) | 1066 | NA | 2.5 |
| *De novo* silent (sibling controls) | 468 | NA | 2.5 |

**Fig 2.**
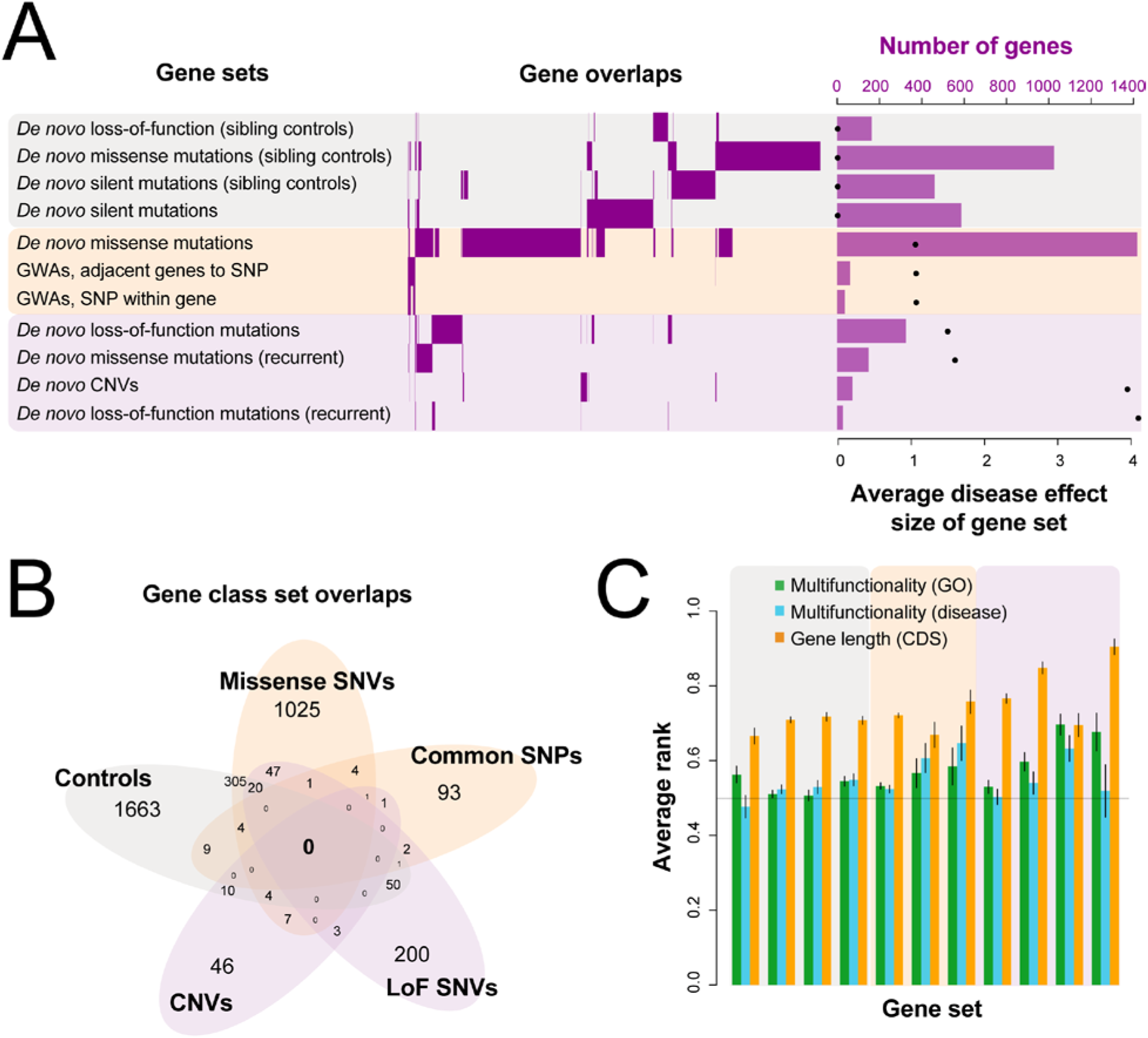
Characterization of autism candidate gene sets. (A) We first classified the 11 gene sets used in the study into three larger groups: no effects, weak effects and strong effects. We see little overlap in the individual gene sets themselves (mid panel). The total number of genes in each set also varies (right-most panel), and is negatively correlated with the average effect size (r_s_=-0.69). (B) Control gene sets overlap significantly with missense genes (333 genes hypergeometric test p=2.54e-6), common SNPs gene sets (11 genes, p=4.5e-3), and the loss-of-function (LoF) SNV gene sets (71 genes, p=3.2e-3) but not the CNV gene sets (14 genes, p=0.03). Missense and common SNPs overlap significantly (4 genes, p=2.4e-4). However, loss-of-function SNVs do not overlap significantly with either common (4 genes, p=0.62), missense (68 genes, p=0.75), or CNVs (3 genes, p=0.37). (C) Common biases that affect studies are gene length and number of functional annotations. The average standardized rank (+/- SE) of genes with respect to these properties shows that the "rare" disease sets contain longer genes but are not much more multifunctional than random.

### Calculating functional convergences

Our functional tests, described below, return p-values which are dependent on the size of the gene set being considered. The statistical tests differ depending on the mode of analysis (e.g., enrichment or network), but by 'functional convergence' we simply mean significance (p-value) after correcting for the set size, typically by downsampling. For the downsampling, we took a subset of genes, recalculated the p-value and then took geometric means of the adjusted p-values. Throughout, where we write 'functional convergence' it is possible to read 'p-value after correcting for set size'.

#### Network connectivity

We measure the clustering of sets of genes within networks through the use of a network modularity calculation. We compare the degree of connections a gene has to all the genes in the network (global node degree), and to those of interest within the sub-network they form (local node degree). The null expectation is that genes will be connected equally well to genes within the sub-network as to those outside. Genes with large positive residuals have more weighted internal connections than external connections, implying a well inter-connected module. We test the significance of this distribution of residuals to a null set (random similarly sized set of genes, Mann-Whitney-Wilcoxon test, wilcox.test in R) to determine our test statistic.

#### Gene set enrichment testing

As a way to determine the level of enrichment of the candidate gene sets within other functional sets, we used a hypergeometric test with multiple test correction (phyper in R). The downsampled p-value was used as the functional convergence measure.

#### Disease gene property testing

For the disease gene scoring properties, we tested the significance of the scores of the candidate genes using the Mann-Whitney-Wilcoxon test (wilcox.test in R). The functional convergence was the p-value of this test.

### Measuring functional convergence trends

For each gene property tested, we then measured the "trend" by calculating the correlation of the ranked functional effect sizes of our gene sets, whereby the gene sets are ordered according to their effect size ranks. A positive correlation is one where the function tested is correlated with our ordering. We computed this using Spearman's rank coefficient to capture the degree of variation, but the significant subsets identified are generally robust to choice of measurement metric such as the Pearson's coefficient. We limited our functional convergence tests to the subset of functions where at least one gene set of the 11 showed a significant functional convergence signal (p<0.05). In essence this filtering removes gene sets where there are, for example, no overlaps with any disease sets and should not affect our analysis.

### Determining significance of the functional convergence trends

To calculate a null, we permute the labels of the gene sets, and calculate the functional convergence trends. Note that in the ranked case, this is simply the null distribution of a spearman correlation, with similarly associated significances. We first filter for functional tests where any one of the disease and control gene sets have a functional convergence of 0.05, but report both pre- and post-filtering results. Because our hypothesis (and test) are concerned with the ordering of functional effect sizes, filtering so that the data has at least one significant value changes the null distribution only slightly (e.g., probability of ties). We calculate the number of significant correlations based on the false discovery rate (FDR) at 0.01 and 0.05. Known confounds of disease gene sets are gene length [59] and gene multifunctionality, and to test this we generated matched gene set controls by sampling genes with similar gene lengths, GO multifunctionality and disease multifunctionality measures. Using the ranked CDS (coding DNA sequence) region of the genes, we generated sets of genes of similar ranked length distributions to the 11 real gene sets in the analysis. Downsampled, we then ran the analyses on these gene sets that are specifically not involved in the phenotype. This was repeated for multifunctionality as calculated using GO and then disease (using Phenocarta [60]).

## Results

### Little overlap of the autism candidate genes across gene sets of different effect sizes

We find genes with loss-of-function *de novo* mutations to be little implicated in GWA studies, with only 4 candidate genes overlapping those two sets (**Fig 2B,** hypergeometric test p=0.76). Interestingly, the more recurrent genes in the loss-of-function *de novo* set, the more unlikely they are to be found in other gene sets. For gene sets with the lower average effect sizes (e.g., the genes with missense mutations), their overlap with other gene sets is greater, in particular with the control sets (**Fig 2B,** hypergeometric tests *p*~4.4e-3 to 2.4e-6). The *de novo* variants are conditioned on being rare (low frequency) and novel by not appearing in the parents. The SNPs used in GWAS are generally conditioned on being common by having minor allele frequencies greater than 0.05 [61]. Even if this filtering is done on the variant level, and not on the gene level, it still creates selection trends within our observations of variants and thus genes. This is possibly a version of Berkson's effect[62] - where selecting for an outcome generates negative correlations between potential causes for it. An additional cause is largely technical; since we've conditioned on frequency, genes with higher mutability are depleted in our rare lists, and enriched in our common lists. Thus the lack of overlap is at least potentially not largely reflective of underlying genetics or biology, but likely due to the selection bias in obtaining them. There is also poor overlap within the rarer variation itself, for instance of genes within CNVs and those with loss-of-function SNVs (3 genes, *p*~0.37); there is generally a discrepancy between study designs focused on (different) sources of rare variation, and not just rare versus common. It should be noted that whether biological or technical, the lack of overlap does nothing to discredit either common or rare variation as a contributor to the disease - but highlights the need for a framework to combine and analyze the results of these studies that is aware of these biases and can distinguish biology from technical effects.

### Functional convergence trends as shown through enrichment and connectivity tests

While enrichment analysis is comparatively straightforward, we demonstrate an example in **Fig 3A** using the genes with *de novo* loss-of-function mutations from lossifov et al,[1] (341 genes) and their overlap with essential genes (see Methods). In **Fig 3A**, we represent this enrichment test as a Venn diagram of the overlap of the candidate disease gene set with the essential gene set, and calculate the significance of the overlap with a hypergeometric test (n=82, *p*~9.8e-9). We continue this analysis on the other candidate disease gene sets from recent ASD studies, varying across study designs and technologies (WES, GWAS and arrays). Splitting each gene set by mutational class, recurrence and gender, we perform the same hypergeometric tests. To make comparable assessments between studies and gene sets, we calculate the functional convergence by downsampling - selecting a subset of genes within that set and averaging the results over a 1000 permutations (schematic in **Fig 3B**). Taking a representative set of studies (**Table 1**), we use the degree of disease effect to rank these sets, noting that recurrence leads to a higher effect size even for variation and study designs of the same class by reducing the number of false positives. Placing the controls sets on the far left, and the highest disease rank set (recurrent *de novo* loss-of-function genes) on the far right, and plotting their functional effect values, we observe an upward trend (**Fig 3C** Spearman's r_s_=0.95, Fisher's transformation p<8.24e-06). The slope (i.e., the correlation) of this trend line represents the "functional convergence trend", with higher correlations indicating higher functional effects.

**Fig 3.**
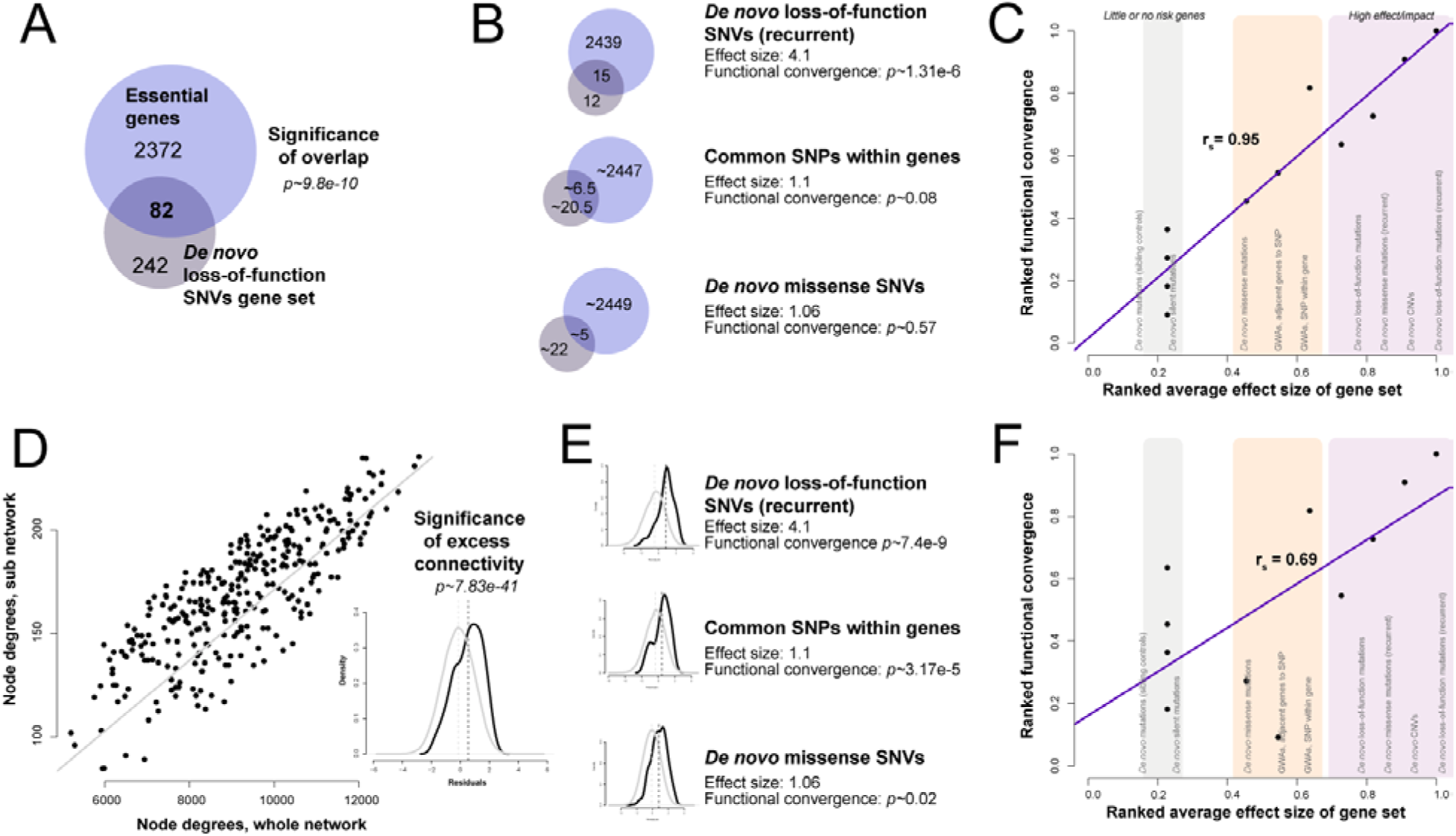
Functional properties of disease gene sets are tested using gene set enrichment (top panels) and coexpression network connectivity (bottom panels). (A) Gene set enrichment is calculated with a hypergeometric test. A large number (34%) of the genes in the *de novo* loss-of-function set overlap with essential genes (hypergeometric test *p*~9.8e-10). (B) This is repeated across all disease gene sets, (a subset shown here). Sample size is controlled through downsampling. Gene sets with the higher effect sizes also have the higher functional convergences. (C) We can now demonstrate how to calculate the functional convergence trend for the "essential genes" test. The disease gene sets are ordered by an estimate of the average effect size of genes within the set (from low to high on the x-axis) and the functional convergence is of that disease gene set is plotted (y-axis). A trend between the effect size of the candidate genes and their essentiality can be clearly observed. The network connectivity functional test (D) consists of calculating the ratio of disease genes' total connectivity (node degrees calculated from the whole network; sum of their connections) to their internal connectivity (node degrees of their subnetwork; sum of their connections to one another). The line (in grey) reflects the expected values if there is no preferential connectivity within the set. We see that a large number (72%) of the genes lie above the identity line. The Wilcoxon p-value of the mean residuals is shown in the inset (*p*~7.83e-41) (E) Once again, controlling for sample size through downsampling, the functional convergence of each gene set is calculated (subset shown). (F) A weak trend between the effect size of the candidate genes and their degree of co-expression is visible. Empirical nulls are calculated by permuting disease gene sets and FDRs through a Benjamini-Hochberg correction against the resultant functional convergence trends.

A less common (likely due to complexity) yet important functional test is network connectivity. Genes that are co-regulated or form parts of a functional unit, protein complex or pathway, are preferentially co-expressed, and this information is captured in co-expression networks. We next demonstrate how network-style effect sizes can be similarly calculated through a modularity analysis. In **Fig 3D**, we plot the global node degrees (x-axis) against their connectivity to the remainder of genes in the set (y-axis). In the null (grey line), the genes would be connected to other autism genes in proportion to the incidence of those genes within the genome. Deviations from this null across all genes generate excess modularity within this set (studentized residuals shown in inset **Fig 3D**) and determine the statistical results reported for the set overall (Wilcoxon test). A large number of genes are highly interconnected in this set, as shown by the number of points above the line (Wilcoxon test on the studentized residuals, *p*~7.83e-41). It is important to note that this network analysis is calculated against the empirical null for each gene individually (x-axis) and so is unaffected by any gene-specific bias (such as length). Only higher-order topological properties across gene-gene relationships for a given gene can produce a signal. Even assortativity, the tendency for genes of high node degree to preferentially interact, is quite low within this data (r=0.064). As in the previous steps, we repeat the network connectivity tests across all gene sets (**Fig 3E**), also downsampling to calculate the functional convergence. Once again, gene sets with higher proportion of burden genes correlate with functional convergence tests (**Fig 3E**, Spearman's r_s_=0.69, Fisher's transformation p<0.02).

### A subset of functional properties are correlated with disease effect sizes

We extend our analysis to other disease gene property tests, and calculate their effect size correlations, plotting the distribution of correlations in **Fig 4A** (4210 functional tests performed, 4164 with calculable correlations). We then calculated the null distribution for the variation across effect sizes by permuting the estimated effect size for each real set and rerunning our analysis. Only limiting our functional tests to those where we had at least one gene set returning a significant enrichment signal, we observe a strong signal (61 tests, **Fig 4A**, 14 functions FDR<0.01 **Table 2**). Reducing the stringency of the underlying enrichment (383 tests, **Fig 4B**), we observe a weaker signal (10 functions FDR<0.01. Removing the underlying enrichment constraint, we observe that most functional tests are ordered consistent with the null, with a few highly correlated functions (**Fig 4C** enrichment at positive end, 3 functions FDR<0.01). The results are broadly reassuring that some weak artifact is not driving the tendency of the functional convergence and effect size to be correlated because that correlation occurs almost exclusively where the underlying tests themselves are detecting significance. In other words, the ordering of significances is only non-random where the underlying values are also non-random. We focus on the 14 functional properties identified in the first filtered assessment (**Table 2**).

**Fig 4.**
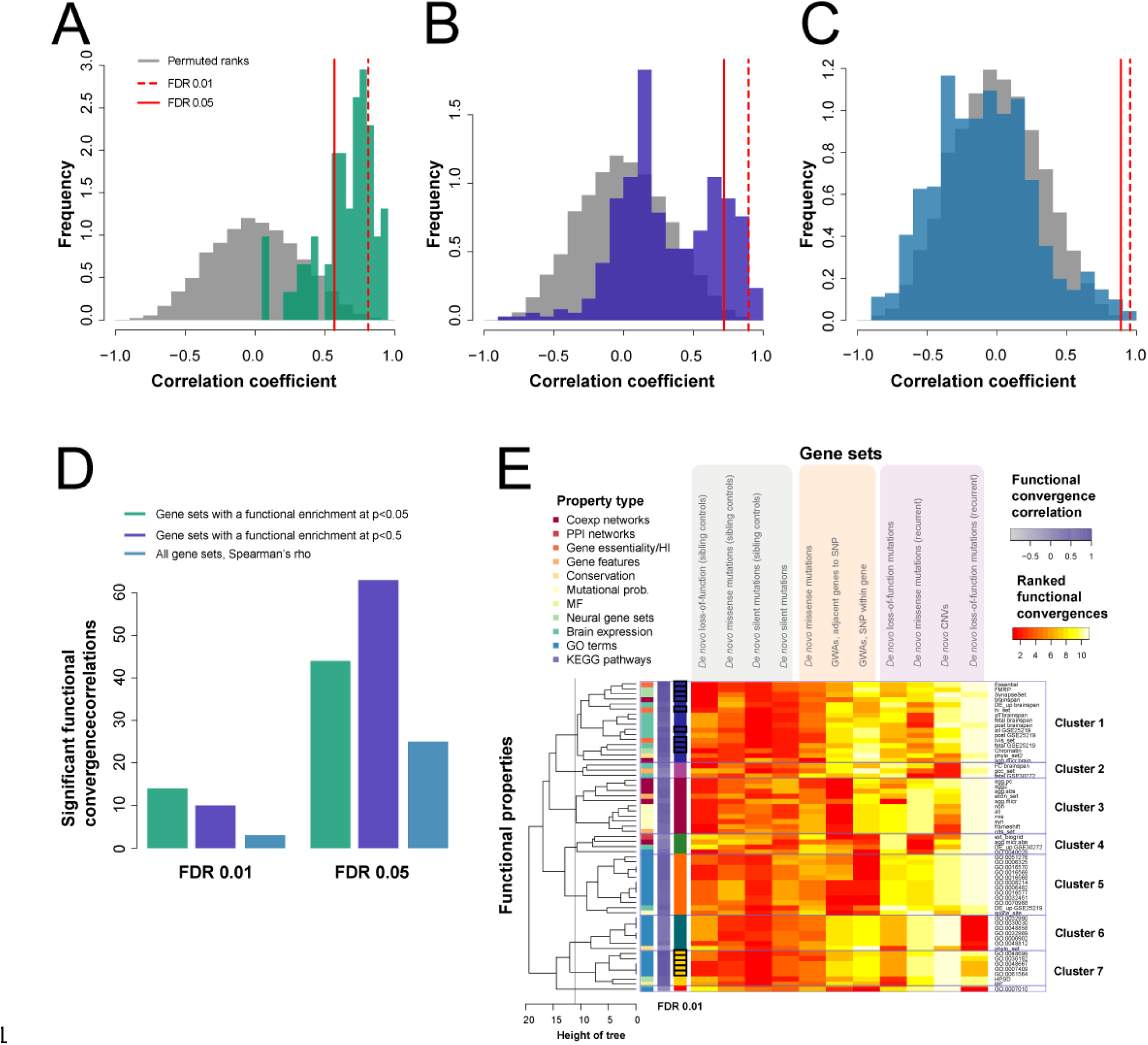
Clustering disease property tests by their functional convergence trends. (A) The correlations of the ranked effect size trends are significantly different from null distributions for the 61 functional properties (effect size permutation null, Student's paired T-test *p* < 2.2e-16).We've drawn the red line to indicate the false discovery rates (FDRs) of 0.01, where 14 functions are significant, and 0.05 where 44 are significant. (B) If we filter for the functions with some weak signal in the underling functional tests (p<0.5), 383 correlations are considered with 10 functions as significant (dark blue, Student's paired T-test *p* < 2.2e-16). Note that we are not filtering with respect to our own functional effect size test, which assesses variation in the underlying functional tests, merely that the underlying tests do return some values. (C) And when we have no constraints (all tests, 4164 shown), 3 pass an FDR of 0.01. (D) We enumerate these in a barplot. (E) A heatmap of all the ranked scores of the test gene sets (columns) for the subset of significant effect 61 properties (rows). The properties clustered into 6 groups when we cut the dendrogram at a height of ~12. Their functional convergence correlations (ranked) show that most high correlations cluster (in clusters 1 and 3). White/yellow is a high rank, red is low. The property type is color coded as described in the figure key. A high correlation is shaded purple, low/negative correlations are grey. Clusters are labeled and colored, with functions FDR<0.01 outlined in black.

**Table 2.** Functional properties with significant functional convergence trends

| **Functional property** | **Spearman's correlation** | **FDR** |
| --- | --- | --- |
| Gene essentiality (Georgi et al.) | 0.95 | 0.001 |
| GO:0048699 (generation of neurons) | 0.92 | 0.003 |
| FMRP interactors (Darnell et al.) | 0.91 | 0.003 |
| Brainspan co-expression network | 0.90 | 0.003 |
| SynapseSet (Lips et al.) | 0.90 | 0.003 |
| Haploinsufficiency scores | 0.89 | 0.003 |
| RVIS | 0.86 | 0.007 |
| GSE25219: Overall expression | 0.84 | 0.008 |
| GO:0007409 (axonogenesis) | 0.84 | 0.006 |
| GO:0030182 (neuron differentiation) | 0.84 | 0.006 |
| GO:0048667 (cell morphogenesis involved in neuron differentiation) | 0.84 | 0.006 |
| GO:0061564 (axon development) | 0.84 | 0.006 |
| Chromatin remodeling gene set (Ronan et al.) | 0.83 | 0.006 |
| GSE25219: Fetal expression | 0.83 | 0.006 |

Each property can be defined by its vector of effect sizes across gene sets and so we can cluster the properties by their Euclidean distance in this space. Taking the 61 properties and highlighting the properties that are significant (FDR 0.01), they split into approximately 7 clusters and a singleton (**Fig 4E**). The interesting clusters are 1 and 7 as they have the highest correlations (as depicted by the dark purple scale), and a stronger significant signal from the *de novo* set (white/yellow in heatmap). Cluster 1, specifically, has the most consistent trends and contains the expression analyses (overexpression and fold change), the gene essentiality scores and some of the neural gene sets. Cluster 3 has the co-expression networks clustered, and the mutational probabilities, but is slightly weaker as the control sets also show some enrichment. Cluster 5 contains most of the GO groups. Cluster 6 has some tests which are functionally enriched in the CNV and missense gene sets but are not significantly enriched for any of the genes in the *de novo* recurrent gene set and are thus not showing a substantially positive functional convergence trend. The clustering speaks to the similarity of some of the tests (i.e., GO groups clustering), but also to a likely neuronal signature across the disease gene sets.

### Significant functional properties are consistent with the autism literature

One of the properties with the highest correlation was network connectivity in the BrainSpan co-expression network; however, all disease gene sets had a significant functional convergence with Brainspan, indicating that in addition to the real signal, there is a background signal affecting even control data. In particular, the signal from the silent recurrent mutations in the probands (functional convergence *p*=7.5e-7) shows that control data subject to only one study design may select genes in a highly non-random pattern. Most top scoring disease properties are consistent with the literature on autism candidates such as average RVIS and haploinsufficiency scores, [63] along with gene length and enrichment for FMRP interactors. RVIS scores are highly enriched in the loss-of-function recurrent set and the CNVs, but not significant in any of the other sets (**Fig 5A**); as with any meta-analysis significance in any one set is not necessary for aggregate significance. Genes with high haploinsufficiency scores - those that cannot maintain normal function with a single copy - are overrepresented in the loss-of-function recurrent genes, and there is also a significant effect in the GWAS results. Many interaction networks and traditional functional categories appear to be poor candidates to determine convergence in disease genes, as they cluster control gene sets and sets of low effects as well as those of disease genes. For instance, the extended PPI network has a high effect in the sibling controls sets (e.g., silent functional convergence *p*~1.3e-5, **Fig 5B**).GO terms and KEGG pathways typically do not survive correcting for multiple testing, although there is a general deviation from the null and the extremal GO functions are concordant with the known literature (e.g., GO: 0016568 chromatin modification hypergeometric test *p*~1e-3 for the *de novo* recurrent set or GO: 0048667 cell morphogenesis involved in neuron differentiation, hypergeometric test *p*~0.04 for the CNV set). So although functional convergence trends are concentrated in more clearly disease related-properties such as RVIS, traditional functional categories from, e.g., GO remain of modest use.

**Fig 5.**
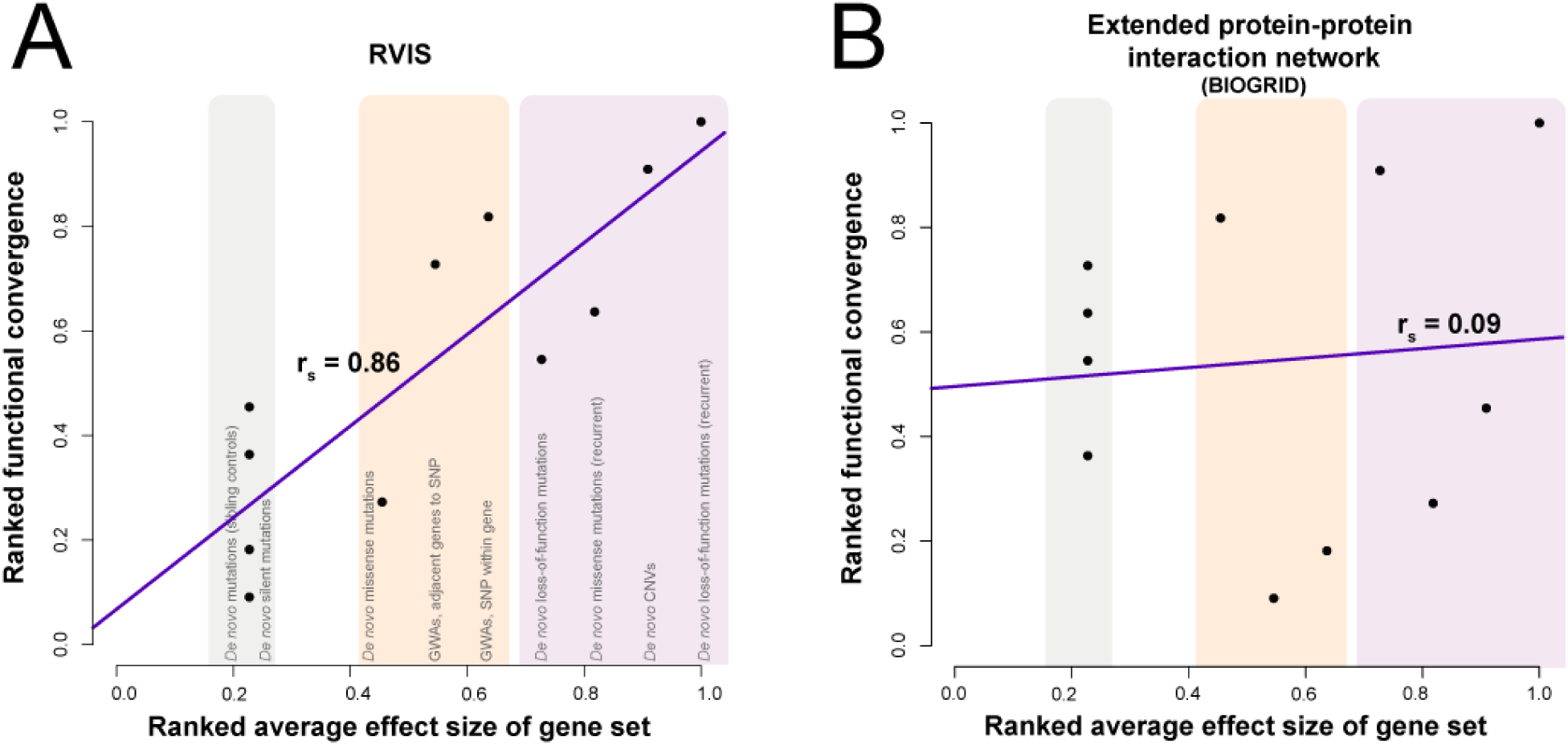
Example correlations and slopes of functional convergence. (A) RVIS enrichment test has a very high correlation across the gene sets (B) while the extended PPIN has mostly artifactual signal as demonstrated by the flatness of the line, highly elevated from the null but in no effect-correlated way and suggestive of consistent biases.

### Robustness and relative contributions of study designs and variants

In order to determine whether the functional convergence trend rose preferentially from a subset of studies, we conducted a series of robustness analyses (**Additional file 1: Fig S1**). Ideally, the significant functional convergence trend we see is due only to effect size estimates across studies which are themselves robust. Nor do we want the trends to be strongly affected by ordering of the gene sets with similar effect sizes. Even though the average effect sizes for the GWAS sets were the same, the number of false positives within these sets varies, and this was incorporated into the ranking scheme. It is also arguable that the silent mutations in the probands may have some regulatory effect, or are false negatives. As a more stringent test, we removed whole classes of variants from the analyses (e.g., all the controls or all the common/weaker gene sets) and calculated the trends once again (**Additional file 1: Fig S1D-F**). This is a negative control experiment in the sense that if the functional convergence trend arises meta-analytically, it should be largely robust to changing things we are not certain about (e.g., as above, whether effect sizes are 1.1 or 1.09) and not robust to changing things we are certain about (common variants play some role in autism). Removing either controls, genes sets with the highest effects or the common variants from the trend analyses removes all the number of significant correlations although some deviation from the null remained (**Additional file 1: Fig S1**). When rare variants are excluded, the distribution of correlations is most similar to the null, but still significantly different (Student's paired T-test *p*~0.03), while the total significance of the test is closest to the full version when common variation is excluded (Student's paired T-test *p*~8.2e-7). Since our common data is likely the weakest due to the tremendous focus of autism data collection toward rare variation in the SSC, this makes sense, but common variation still contributes substantial joint signal. These tests confirm that the approximate order of gene sets by effect sizes correctly drives the results and that we are robust to minor variation in the exact effect sizes listed, but do rely on the joint use of the extremely divergent study results (rare and common) within the meta-analysis to attain significant results.

To control for the impact of gene length and multifunctionality (number of functions a gene is listed as possessing), we repeated a control version of our analyses. In this case, the real disease gene sets were swapped out with gene sets matched with respect to multifunctionality or length. We then reran the evaluation of functional convergence trends to determine if any previously identified properties arise as correlated with these control sets (ordered by their match to a specific disease sets, e.g., gene length distribution). Repeating the analysis in this control case, we find the derived correlations are for the most part extremely similar to the null (reference). We can additionally use these controls versions as a slightly more stringent null distribution for expected correlations when we evaluate the real disease sets. In the analysis where we do not condition on the underling tests having reached some level of significance (as in **Fig 4A**), we see even more correlations passing significance (**Additional file 1: Fig S2**), indicating the multifunctionality or gene length do little to explain the general trends we see.

Promiscuous or absent enrichment have both historically been problematic within disease gene data; both diminish the specificity of functional results. When too many functions are returned from an analysis, we need to cherry pick and with too few, we have no "leads" and are left in the dark. We suggest that the strong aggregate effect we see and small number of significant functions is likely near to a useful and biologically plausible type of specificity for downstream analysis, as suggested by the fact that ad hoc filtering (i.e., top ten lists) usually are at about this level when not constrained by significance. Our set of functional tests and results are shown in **Additional file 2: Table S1** and the full data set is available online.

One potential failure mode of this analysis comes from the GWAs we have used. Because the number of autism GWAs available and well-powered for analysis was relatively small, we used a combined psychiatric genomics dataset, which included bipolar and schizophrenia. We now wish to test how specific our results were to our disease and not a signal of GWAs in general. We repeated our analysis using each of the 148 GWAS traits in the GWAS catalog that had enough genes to be included in our tests. We did not recalculate the effect sizes specifically for each, but used the mean estimates from the autism set. Using the number of correlations calculated as significant to rank the 148 traits, the top ten traits include the autism and schizophrenia GWAs, and a few larger studies such as "Body mass index" (**Additional file 1: Fig S3**). This is a fairly striking confirmation of our original hypothesis: the degree of correlation between functional convergence is so specific that it correctly distinguishes particular disease sets as belonging to the same trend (as defined by a particular disease). The larger GWAs, also found in the top 10, are not related to psychiatric disorders show a signal in very broad disease properties, such as the gene mutability scores.

### Expanding the functional gene tests show no further significant properties

We wished to see if we could find other significant associations if we expanded our repertoire of functions within each type of test. Our first set of network analyses focused on general aggregate co-expression networks and brain sample only aggregates. In most analyses, researchers use individual datasets to build their networks and we wished to compare our results to these. Thus we expanded our tests to a total of 540 networks. We repeated the same analysis, using an additional 6 PPI networks, 76 condition specific networks (tissues, sex, and age), and a further 454 RNAseq co-expression networks (227 across 18K protein coding genes and 227 across 30K transcripts). Once again, we see functional convergence across almost all the gene sets with little ordered trend by effect size. The network convergence exists in even the control data and is therefore likely due to study selection biases alone; none pass an FDR of 0.01 (**Additional file 1: Fig S4A**).

Initially, we focused on expression data for the brain, but were curious about how tissue-specific these patterns were, or whether the genes were generally highly expressed. To this end, we repeated the expression analyses using tissue specific expression datasets from GTEx data [31]. We were also curious to determine if we could see sex specific differences, and used additional data from the GEUVADIS project [64]. Repeating the functional convergence trends on all these expression datasets shows little to no significant expression in the individual gene sets, and no significant functional correlations (**Additional file 1: Fig S4B**). The functional test with the greatest correlation were also from brain specific expression datasets (r_s_=0.78).

One last set of gene properties typically used by researchers in their analyses are the curated gene sets from MSigDB. We repeated our analyses on all 8 collections, and calculated the functional convergence trends, using the hypergeometric test as in the case of calculating enrichment in GO. The gene sets range from curated data sets from the known literature, to computationally derived gene sets from cancer microarrays. Perhaps unsurprisingly, we see no enrichment in these gene sets (**Additional file 1: Fig S5**), as most are inflammatory or oncogenic collections, or versions of GO terms and KEGG pathways which we had already found to have no enrichment.

## Discussion

Our contribution in this work has been to establish that there is a significant correlation between the effect size of candidate autism genes and the degree to which tests assessing their functional convergence find a signal. As with any meta-analysis, the hope is that by incorporating multiple data, the aggregate signal may be stronger. While our work suggests an approach to do this and shows strong statistical trends, we anticipate that divisions in the field of genetics may play some role in the interpretation of this work [65, 66]. GWAS researchers may question the power of our GWAS analysis and assume that if it just had a large enough ‘N’, it alone would be the dominant player in understanding psychiatric disease. Similarly, they may suspect rare variants of reflecting ‘anomalous’ versions of the disorder and thus think they are less likely to be specifically linked to autism. Rare variant researchers may question the precision of the data underlying our rare variant analysis and assume that if we just had large enough ‘N’ to remove false positives, it would be the dominant player in understanding psychiatric disease. Similarly, they may suspect that GWAS data is affected by confounds such as population stratification. Both reactions are perfectly reasonable. Our analysis does not establish that all functional properties are distributed across all classes of autism, but rather, for a subset, there is a very significant trend. This is further supported by the functions that arise being either of specific relevance to autism or of well-known importance to disease in general; however, even this division implies that we are capturing multiple factors affecting the genetic architecture of disease.

Likewise, our specific experimental design for assessing a relationship between functional convergence and effect size may well be open to elaboration and emendation. Our principle interest was in ensuring that any observed trends would be reflective of tests and data used within the literature and not choices of our own. Thus, we aimed for the simplest and most conventional means of assessing functional convergence and focused on being exhaustive (in terms of properties) within this domain, rather than optimizing our design for observing functional convergence trends. We also developed a framework which is readily extensible to new tests, regardless of their form or complexity. In each case, the test can be equivalently applied to the downsampled disease gene sets and the significance of any correlation simply calculated as would be conventional via permutation test. This is both simple, general, and allows easy comparison across studies using the equivalent approach; however, more theoretically grounded or alternative means of calculating functional convergence trends are surely possible. Particular weaknesses in our design are our use of non-parametric tests and downsampling to control for set size. These are, we think, natural choices for robustness but more finely tuned alternatives are likely to exist and could easily be a target of research since our results suggests the observation of key functional convergence trends is highly robust and salient within the data.

As the number of disease gene sets expands, and further refinement of risk assessment is achieved, the resolution of functional convergence trends should grow. Indeed, incorporating effect size as a metaanalytic constraint offers a diverse range of novel applications. That integration may be across study designs and classes of variation, as we have done, or may involve phenotype or other properties. So, for example, one could determine functional convergence trends that grow or shrink depending on how patients were classified, or even broken down in a sex-specific manner for interpreting protective effects. More broadly, as data and the means for obtaining it grows, techniques to statistically assess its structured dependencies will grow more useful and important. Our robustness analysis speaks to this in that while we are robust to modest losses of data, it is clear that more data will only improve the signals of the individual classes. More finely-tuned effect size estimates and better separations of the gene sets and variant classifications will also help refine the distinction between biological and artifactual signals, ideally allowing us conduct yet more focused study designs in a productive feedback loop.

## Conclusions

In this work we have found that the stronger the effect size of autism candidate genes, the more likely they are to exhibit a joint functional signal. The functional properties identified exhibit some specificity to autism and neuropsychiatric disease (e.g. FMRP interactors), but also some more general links to disease (e.g., RVIS). While there remains substantial heterogeneity between study designs and the genetic architectures of disease which they may uncover, we have shown that there is some commonality across study designs. The commonality across study designs is not a literal overlap in risk genes, or even functional effect, but that functions weakly identified in GWA studies are likely to be more strongly identified in rare variation studies. As evidence for autism and other disorders continues to develop and continues to be heterogeneous with respect to ascertainment biases and study designs, we suspect approaches related to the one we describe will be of increasing importance.

## Abbreviations

ASD: autism spectrum disorder
CNV: copy number variant
FDR: false discovery rate
FMRP: fragile X mental retardation protein
FWER: family-wise error rate
GO: Gene Ontology
GWAS: genome-wide association studies
HI: happloinsufficiency
LoF: Loss-of-function mutations
PGC: Psychiatric Genomics Consortium
PPI: protein-protein interactions
PPIN: protein-protein interaction network
RVIS: Residual Variation Intolerance Score
SNP: single nucleotide polymorphism
SNV: single nucleotide variant
SSC: Simons Simplex Collection
WES: whole exome sequencing
WGS: whole genome sequencing

## Availability of data and materials

The datasets supporting the conclusions of this article are included within the article and in the additional files.

## Additional files

Additional file 1: Figures

Additional file 1: Fig S1 Trend line robustness analysis.

Additional file 1: Fig S2 Functional convergences null for matched length and multifunctionality controls

Additional file 1: Fig S3 Functional convergence correlation/trend distributions for GWA studies.

Additional file 1: Fig S4 Functional convergence correlation/trend distributions for all network connectivity tests and all gene expression tests

Additional file 1: Fig S5 Functional convergence correlation/trend distributions for all MSigDB collections.

Additional file 2: Tables

Additional file 2: Table S1 Functional convergence correlation/trend distributions

Additional file 2: Table S2 Additional expression functional convergences and correlations/trends

Additional file 2: Table S3 Additional gene properties functional convergences and correlations/trends

Additional file 2: Table S4 Additional MSigDB functional convergences and correlations/trends

Additional file 2: Table S5 Additional network connectivity functional convergences and correlations/trends

## Authors’ contributions

SB wrote the manuscript, and conducted the experiments. JG wrote the manuscript and designed the experiments. All authors read and approved the final manuscript.

## Acknowledgments

We would like to thank members of the CSHL Wigler lab for access to their data. We thank Paul Pavlidis for helpful comments on a draft of the manuscript.

## References

1. SebatJ, LakshmiB, MalhotraD, TrogeJ, Lese-MartinC, WalshT, YamromB, YoonS, KrasnitzA, KendallJ, et al: Strong Association of De Novo CopyNumber Mutations with Autism. Science 2007, 316:445–449.

2. ChristianSL, BruneCW, Sudi J, KumarRA, LiuS, KaraMohamedS, BadnerJA, MatsuiS, ConroyJ, McQuaidD, et al: Novel Submicroscopic chromosomal abnormalitiesdetected in Autism Spectrum Disorder. Biological psychiatry 2008, 63:1111–1117.

3. MarshallCR, NoorA, VincentJB, LionelAC, FeukL, SkaugJ, ShagoM, MoessnerR, PintoD, RenY, et al: Structural Variation of Chromosomes in Autism SpectrumDisorder. American Journal of HumanGenetics 2008, 82:477–488.

4. GlessnerJT, WangK, CaiG, KorvatskaO, KimCE, WoodS, ZhangH, EstesA, BruneCW, BradfieldJP, et al: Autism genome-wide copy number variation reveals ubiquitin and neuronal genes. Nature 2009, 459:569–573.

5. MaDQ, SalyakinaD, JaworskiJM, KonidariI, WhiteheadPL, AndersenAN, HoffmanJD, SliferSH, HedgesDJ, CukierHN, et al: A genome-wide association study of autism reveals a common novel risk locus at 5p14.1. Annals of humangenetics 2009, 73:263–273.

6. WelterD, MacArthur J, Morales J, Burdett T, Hall P, Junkins H, Klemm A, Flicek P, Manolio T, Hindorff L, Parkinson H: The NHGRI GWAS Catalog, acurated resource of SNP-trait associations. Nucleic Acids Research 2014, 42:D1001–D1006.

7. Subramanian A, Tamayo P, Mootha VK, Mukherjee S, Ebert BL, Gillette MA, Paulovich A, Pomeroy SL, Golub TR, Lander ES, Mesirov JP: Gene set enrichment analysis: A knowledge-based approach for interpretinggenome-wide expression profiles. Proceedingsof the National Academy of Sciences 2005, 102:15545–15550.

8. Zoubarev A, Hamer KM, Keshav KD, McCarthy EL, Santos JRC, Van Rossum T, McDonald C, Hall A, Wan X, Lim R: Gemma: a resource for the reuse, sharing and meta-analysis of expression profiling data. Bioinformatics 2012, 28:2272–2273.

9. O'Roak BJ, Deriziotis P, Lee C, Vives L, Schwartz JJ, Girirajan S, Karakoc E, MacKenzie AP, Ng SB, Baker C: Exome sequencing in sporadic autism spectrum disorders identifies severe de novomutations. Naturegenetics 2011, 43:585–589.

10. Sanders SJ, Murtha MT, Gupta AR, Murdoch JD, Raubeson MJ, Willsey AJ, Ercan-Sencicek AG, DiLullo NM, Parikshak NN, Stein JL: De novo mutations revealed by whole-exome sequencing are strongly associated withautism. Nature 2012, 485:237–241.

11. Iossifov I, Ronemus M, Levy D, Wang Z, Hakker I, Rosenbaum J, Yamrom B, Lee Y-h, Narzisi G, Leotta A, et al: De Novo Gene Disruptions in Children on the Autistic Spectrum. Neuron 2012, 74:285–299.

12. Neale BM, Kou Y, Liu L, Ma'ayan A, Samocha KE, Sabo A, Lin CF, Stevens C, Wang LS, Makarov V, et al: Patterns and rates of exonic de novo mutations in autism spectrumdisorders. Nature 2012, 485:242–245.

13. Lohmueller KE, Pearce CL, Pike M, Lander ES, Hirschhorn JN: Meta-analysis ofgenetic association studies supports a contribution of common variants to susceptibility to common disease. Nat Genet 2003, 33:177–182.

14. Wang K, Zhang H, Ma D, Bucan M, Glessner JT, Abrahams BS, Salyakina D, Imielinski M, Bradfield JP, Sleiman PM, et al: Common genetic variants on 5p14.1 associate with autism spectrumdisorders. Nature 2009, 459:528–533.

15. Cobb JP, Mindrinos MN, Miller-Graziano C, Calvano SE, Baker HV, Xiao W, Laudanski K, Brownstein BH, Elson CM, Hayden DL, et al: Application of genome-wide expression analysis to human health and disease. Proceedings of the National Academy of Sciences of the United States ofAmerica 2005, 102:4801–4806.

16. Deng Q, Ramsköld D, Reinius B, Sandberg R: Single-Cell RNA-Seq RevealsDynamic, Random Monoallelic Gene Expression in Mammalian Cells. Science 2014, 343:193–196.

17. Bauer-Mehren A, BundschusM, Rautschka M, Mayer MA, Sanz F, Furlong LI: Gene-Disease Network Analysis Reveals FunctionalModules in Mendelian, Complex and EnvironmentalDiseases. PLoS ONE 2011, 6:e20284.

18. Ben-David E, Shifman S: Networks of Neuronal Genes Affected by Common and RareVariants in Autism Spectrum Disorders. PLoS Genet 2012, 8:e1002556.

19. Sakai Y, Shaw CA, Dawson BC, Dugas DV, Al-Mohtaseb Z, Hill DE, Zoghbi HY: Protein Interactome Reveals Converging MolecularPathways Among Autism Disorders. ScienceTranslational Medicine 2011, 3:86ra49.

20. Sullivan PF, Daly MJ, O'Donovan M: Genetic architectures ofpsychiatric disorders: the emerging picture and itsimplications. Nat RevGenet 2012, 13:537–551.

21. Schizophrenia WorkingGroup of the Psychiatric Genomics C: Biologicalinsights from 108 schizophrenia-associated geneticloci. Nature 2014, 511:421–427.

22. Miles JH: Autism spectrum disorders--a geneticsreview. Genetics inMedicine 2011, 13:278–294.

23. Cross-Disorder Group ofthe Psychiatric Genomics C: Identification of riskloci with shared effects on five major psychiatric disorders: agenome-wide analysis. Lancet 2013, 381:1371–1379.

24. Gibson G: Rare and common variants: twentyarguments. Nat RevGenet 2012, 13:135–145.

25. lossifov I, O'Roak BJ, Sanders SJ, Ronemus M, Krumm N, Levy D, Stessman HA, Witherspoon K, Vives L, Patterson KE, et al: The contribution of de novo coding mutations to autism spectrum disorder. Nature 2014, 515:216–221.

26. Levy D, Ronemus M, Yamrom B, Lee YH, Leotta A, Kendall J, Marks S, Lakshmi B, Pai D, Ye K, et al: Rare de novo and transmitted copy-number variation in autistic spectrum disorders. Neuron 2011, 70:886–897.

27. Willsey AJ, Sanders Stephan J, Li M, Dong S, Tebbenkamp Andrew T, Muhle Rebecca A, Reilly Steven K, Lin L, Fertuzinhos S, Miller Jeremy A, et al: Coexpression Networks Implicate Human Midfetal Deep Cortical Projection Neurons in the Pathogenesis of Autism. Cell 2013, 155:997–1007.

28. Sugathan A, Biagioli M, Golzio C, Erdin S, Blumenthal I, Manavalan P, Ragavendran A, Brand H, Lucente D, Miles J, et al: CHD8 regulates neurodevelopmental pathways associated with autism spectrum disorder in neural progenitors. Proceedings of the National Academy of Sciences 2014, 111:E4468–E4477.

29. Hormozdiari F, Penn O, Borenstein E, Eichler EE: The discovery of integrated gene networks for autism and related disorders. Genome Research 2015, 25:142–154.

30. Verleyen W, Ballouz S, Gillis J: Positive and negative forms of replicability in gene network analysis. Bioinformatics 2015.

31. Lonsdale J, Thomas J, Salvatore M, Phillips R, Lo E, Shad S, Hasz R, Walters G, Garcia F, Young N, et al: The Genotype-Tissue Expression (GTEx) project. NatureGenetics 2013, 45:580–585.

32. Harrow J, Frankish A, Gonzalez JM, Tapanari E, Diekhans M, Kokocinski F, Aken BL, Barrell D, Zadissa A, Searle S, et al: GENCODE:the reference human genome annotation for The ENCODEProject. GenomeResearch 2012, 22:1760–1774.

33. Stark C, Breitkreutz B-J, Reguly T, Boucher L, Breitkreutz A, Tyers M: BioGRID:a general repository for interaction datasets. Nucleic Acids Research 2006, 34:D535–D539.

34. Chua HN, Sung W-K, Wong L: Exploiting indirect neighbours and topological weight to predict protein function from protein-protein interactions. Bioinformatics 2006, 22:1623–1630.

35. Gillis J, Pavlidis P: The role of indirect connections in gene networks in predicting function. Bioinformatics 2011, 27:1860–1866.

36. Brown KR, Jurisica I: Unequal evolutionary conservation of human protein interactions in interologous networks. GenomeBiology 2007, 8:1–11.

37. Peri S, Navarro JD, Kristiansen TZ, Amanchy R, Surendranath V, Muthusamy B, Gandhi TKB, Chandrika KN, Deshpande N, Suresh S, et al: Human protein reference database as a discovery resource for proteomics. Nucleic Acids Research 2004, 32:D497–D501.

38. Schaefer MH, Fontaine J-F, Vinayagam A, Porras P, Wanker EE, Andrade-Navarro MA: HIPPIE: Integrating Protein Interaction Networks with Experiment Based Quality Scores. PLoSONE 2012, 7:e31826.

39. Orchard S, Ammari M, Aranda B, Breuza L, Briganti L, Broackes-Carter F, Campbell NH, Chavali G, Chen C, del-Toro N, et al: The MIntAct project—IntAct as a common curation platform for 11molecular interaction databases. NucleicAcids Research 2014, 42:D358–D363.

40. Rolland T, Ta§an M, Charloteaux B, Pevzner Samuel J, Zhong Q, Sahni N, Yi S, Lemmens I, Fontanillo C, Mosca R, et al: AProteome-Scale Map of the Human Interactome Network. Cell, 159:1212–1226.

41. Szklarczyk D, Franceschini A, Wyder S, Forslund K, Heller D, Huerta-Cepas J, Simonovic M, Roth A, Santos A, Tsafou KP, et al: STRINGv10: protein-protein interaction networks, integrated over the treeof life. Nucleic AcidsResearch 2015, 43:D447–D452.

42. McDowall MD, Scott MS, Barton GJ: PIPs: human protein-protein interactionprediction database. Nucleic AcidsResearch 2009, 37:D651–D656.

43. Blohm P, Frishman G, Smialowski P, Goebels F, Wachinger B, Ruepp A, Frishman D: Negatome 2.0: a database of non-interacting proteins derived by literature mining, manual annotation and protein structureanalysis. Nucleic AcidsResearch 2014, 42:D396–D400.

44. Ronan JL, Wu W, Crabtree GR: From neural development to cognition: unexpectedroles for chromatin. Nature reviewsGenetics 2013, 14:347–359.

45. Bayes A, Collins MO, Croning MDR, van de Lagemaat LN, Choudhary JS, Grant SGN: Comparative Study of Human and Mouse Postsynaptic Proteomes Finds High Compositional Conservation and AbundanceDifferences for Key Synaptic Proteins. PLoSONE 2012, 7:e46683.

46. Collins MO, Husi H, Yu L, Brandon JM, Anderson CNG, Blackstock WP, Choudhary JS, Grant SGN: Molecular characterization and comparison of the components and multiprotein complexes in the postsynapticproteome. Journal ofneurochemistry 2006, 97 Suppl1:16–23.

47. Petrovski S, Wang Q, Heinzen EL, Allen AS, Goldstein DB: Genic Intoleranceto Functional Variation and the Interpretation of PersonalGenomes. PLoSGenet 2013, 9:e1003709.

48. Georgi B, Voight BF, Bucan M: From Mouse to Human: Evolutionary Genomics Analysisof Human Orthologs of Essential Genes. PLoS Genet 2013, 9:e1003484.

49. Huang N, Lee I, Marcotte EM, Hurles ME: Characterising and PredictingHaploinsufficiency in the Human Genome. PLoS Genet 2010, 6:e1001154.

50. Verleyen W, Ballouz S, Gillis J: Measuring the wisdom of the crowds innetwork-based gene function inference. Bioinformatics 2015, 31:745–752.

51. Kanehisa M, Goto S: KEGG: kyoto encyclopedia of genes andgenomes. Nucleic AcidsResearch 2000, 28:27–30.

52. Gillis J, Pavlidis P: The Impact of Multifunctional Genes on "Guilt byAssociation" Analysis. PLoSONE 2011, 6:e17258.

53. Darnell JC, Van Driesche SJ, Zhang C, Hung KYS, Mele A, Fraser CE, Stone EF, Chen C, Fak JJ, Chi SW, et al: FMRP stalls ribosomal translocation on mRNAs linked to synaptic function andautism. Cell 2011, 146:247–261.

54. Lips ES, Cornelisse LN, Toonen RF, Min JL, Hultman CM, Holmans PA, O'Donovan MC, Purcell SM, Smit AB, Verhage M, et al: Functional gene group analysis identifies synapticgene groups as risk factor for schizophrenia. Mol Psychiatry 2012, 17:996–1006.

55. Lek M, Karczewski K, Minikel E, Samocha K, Banks E, Fennell T, O'Donnell-Luria A, Ware J, Hill A, Cummings B: Analysis of protein-coding genetic variation in 60,706 humans. BioRxiv 2016:030338.

56. Levy D, Ronemus M, Yamrom B, Lee Y-h, Leotta A, Kendall J, Marks S, Lakshmi B, Pai D, Ye K, et al: Rare De Novo and Transmitted Copy-Number Variation in Autistic SpectrumDisorders. Neuron 2011, 70:886–897.

57. Colantuoni C, Lipska BK, Ye T, Hyde TM, Tao R, Leek JT, Colantuoni EA, Elkahloun AG, Herman MM, Weinberger DR, Kleinman JE: Temporal dynamics and genetic control of transcription in the human prefrontalcortex. Nature 2011, 478:519–523.

58. Chang J, Gilman SR, Chiang AH, Sanders SJ, Vitkup D: Genotype to phenotyperelationships in autism spectrum disorders. Nat Neurosci 2015, 18:191–198.

59. Ouwenga RL, Dougherty J: Fmrp targets or not: long, highly brain-expressedgenes tend to be implicated in autism and braindisorders. MolecularAutism 2015, 6:16.

60. Portales-Casamar E, Ch'ng C, Lui F, St-Georges N, Zoubarev A, Lai A, Lee M, Kwok C, Kwok W, Tseng L, Pavlidis P: Neurocarta:aggregating and sharing disease-gene relations for theneurosciences. BMCGenomics 2013, 14:129.

61. Bush WS, Moore JH: Chapter 11: Genome-Wide AssociationStudies. PLoS ComputBiol 2012, 8:e1002822.

62. Westreich D: Berkson's bias, selection bias, and missingdata. Epidemiology (Cambridge, Mass) 2012, 23:159–164.

63. Ji X, Kember RL, Brown CD, Bućan M: Increased burden of deleterious variants inessential genes in autism spectrum disorder. Proceedings of the National Academy ofSciences 2016, 113:15054–15059.

64. Lappalainen T, Sammeth M, Friedlander MR, t Hoen PAC, Monlong J, Rivas MA, Gonzalez-Porta M, Kurbatova N, Griebel T, Ferreira PG, et al: Transcriptome and genome sequencing uncoversfunctional variation in humans. Nature 2013, 501:506–511.

65. Krumm N, Turner TN, Baker C, Vives L, Mohajeri K, Witherspoon K, Raja A, Coe BP, Stessman HA, He Z-X: Excess of rare, inherited truncating mutationsin autism. Naturegenetics 2015, 47:582–588.

66. Pinto D, Delaby E, Merico D, Barbosa M, Merikangas A, Klei L, Thiruvahindrapuram B, Xu X, Ziman R, Wang Z, et al: Convergence of Genesand Cellular Pathways Dysregulated in Autism SpectrumDisorders. The American Journal of HumanGenetics 2014, 94:677–694.

